# Visual consequences of saccades explain early cortical response dynamics during natural vision

**DOI:** 10.64898/2026.06.12.731798

**Authors:** Richard Schweitzer, Olaf Dimigen, Christoph Huber-Huber

## Abstract

During natural vision, each saccadic eye movement evokes a large cortical potential known as the lambda response. Traditionally, lambda responses have been considered feedforward responses to new visual input arriving at fixation. In contrast, here we show that they are best explained by the retinal shifts induced by the saccade itself. We first recorded EEG data for thousands of saccades during free viewing of natural scenes and then replayed the same retinal image shifts on a high-speed display during fixation. Temporal response dynamics were virtually identical for real saccades and replayed motion, and most strongly aligned to saccadic peak velocity. A computational model based only on saccade trajectories, 1/f natural-scene statistics, and known human spatiotemporal sensitivity parsimoniously reproduced these dynamics. Our results show that saccade intervals should not be viewed as a gap in visual processing, but as a structured sensory event that shapes human cortical activity during natural viewing.

## Introduction

In natural environments, humans incessantly make eye movements. About three to four times per second, their gaze shifts in rapid step-like movements, so-called saccades, to explore the visual scene in front of them. Being one of the fastest and most frequent of human movements, saccades have drastic visual consequences: They cause high-speed shifts of the entire visual image across the retina. Only during the fixation period after each saccade, the visual system can sample somewhat stable input from the new gaze location. This cyclic process of active vision triggers brain responses denoted as lambda responses. Traditionally recorded with electroencephalography (EEG), they typically peak around 50-100 ms after saccade offsets at occipital sensors. First reported in the 1950s (Evans, 1953; Gastaut, 1951), lambda responses have been the subject of numerous studies. There is ample evidence that they reflect the feedforward sweep of information along the retinotopically organized visual hierarchy, as they vary with the features of the visual stimulus (Gaarder, Krauskopf, Graf, Kropfl, & Armington, 1964; Ries, Slayback, & Touryan, 2018) and remain absent in total darkness, with closed eyes, during steady fixation, or when viewing a uniform visual field (Barlow & Cigánek, 1969; Billings, 1989). Considering these features together with the frequency of saccades in everyday life, lambda responses are among the brain’s most prominent event-related responses under natural conditions.

While it is highly plausible that lambda responses are evoked by visual processing, it still remains largely unknown how exactly they are generated. Previous studies have attributed lambda responses to the superposition of pre- and post-saccadic visual responses (Thickbroom, Knezevič, Carroll, & Mastaglia, 1991; Yagi, 1979), as well as the time course of saccadic suppression (Billings, 1989). More recent studies proposed another mechanism, that pre-saccadic preparatory and predictive processes specifically related to active vision drive the dynamics of lambda responses (Amme et al., 2024; Nolte, Schmidt, Grasso-Cladera, & König, 2025). This idea was supported by the intriguing phenomenon that lambda responses lock more closely to saccade onset than saccade offset. Indeed, when measured relative to saccade offset, the latency of lambda responses decreases as the size of inducing saccades increases (Amme et al., 2024; Dimigen, Sommer, Hohlfeld, Jacobs, & Kliegl, 2011; Gordon, Dalangin, & Touryan, 2024; Kaunitz et al., 2014; Nolte et al., 2025; Yagi, 1979). This finding appears to be at odds with the notion of bottom-up, feedforward visual processing and raises the question how well lambda responses are understood at the mechanistic level.

Here, we provide a principled explanation for the dynamics of lambda responses, demonstrating that they result from the visual transients generated by saccades. Crucially, using modern display technology with high spatial and temporal resolution, we contrasted active exploration of natural scenes with passive viewing where the exact visual consequences of saccades were replayed to fixating observers, revealing that lambda-response latency decreased with increasing saccade size regardless of whether saccades were actively executed. Furthermore, during both active and passive viewing, lambda responses locked to neither saccade onset nor offset but to the intra-saccadic time of saccadic peak velocity. With the help of computational modeling of early visual processing (Schweitzer, Seel, Raisch, & Rolfs, 2025), we derive that lambda responses can be parsimoniously explained by the spatiotemporal dynamics of visual shifts introduced by saccades during natural viewing, without requiring predictive or extra-retinal mechanisms specific to active vision.

## Results

Across a total of four separate experimental sessions, twelve human observers performed a simple object-reporting task (Figure 1A) while EEG and video-based eye-tracking (ET) measurements were conducted. In the first two sessions, observers freely explored a set of 1980 color images retrieved from a widely used natural-scene database (Allen et al., 2022, see Stimuli for details) and, with a probability of 1/8 on each trial, were prompted to indicate whether a certain object category was either present or absent in the most recent scene. The purpose of this viewing condition, henceforth *Saccade* condition, was to record a large number of free-viewing saccades and to measure lambda responses during natural active vision. In the third and fourth sessions, the critical *Replay* viewing condition was presented, in which – during fixation – the same scenes rapidly moved across the screen simulating the previously recorded eye-movement trajectories for a given scene and observer (see Replay trajectories). Importantly, reproducing motion at saccadic speeds requires presentation systems with high temporal resolution (e.g., Duyck, Wexler, Castet, & Collins, 2018; Schweitzer & Rolfs, 2020), which is why we applied a modern OLED monitor capable of variable frame rates up to 480 Hz (Figure 1B; see also Figure S1 for timing evaluations). A total of 143888 pairs of real and replayed saccades entered the analysis, a data set large enough to perform a highly powered, direct comparison of lambda responses with moving and fixating eyes to isolate potential mechanisms specific to active vision.

**Figure 1.**
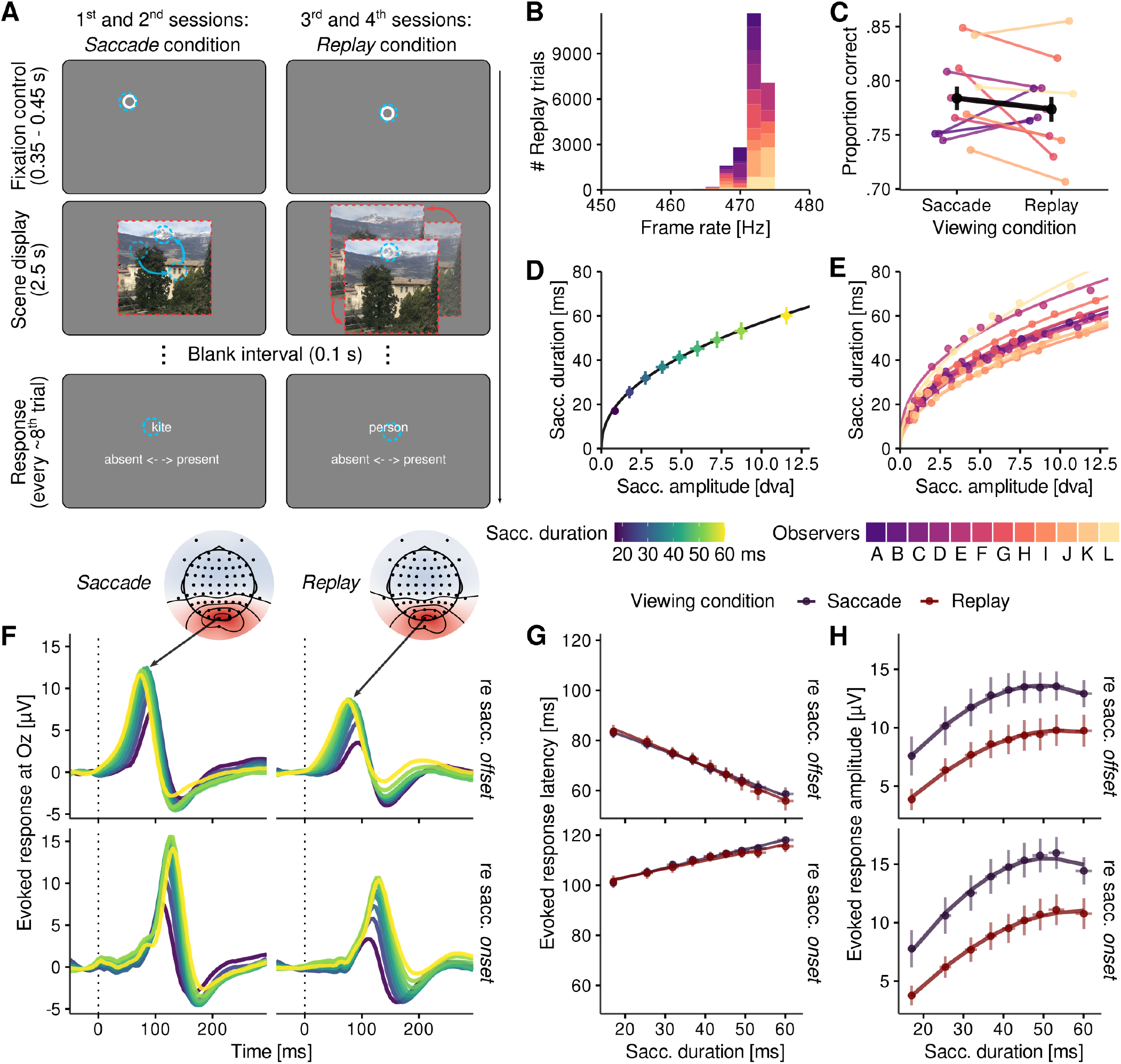
Visually evoked responses to real and replayed saccades. **A** Illustration of experimental paradigm (for details, see Procedure) using active visual exploration in Saccade sessions (left column) and a replay of the visual consequences of that exploration during fixation in Replay sessions (right column). **B** Distribution of timestamp-based measurements of monitor frame rates during Replay trials confirming an empirical frame rate between 470 and 475 Hz. **C** Proportion of correct responses during Saccade and Replay trials. Colors indicate individual observers, the group average is plotted in black. **D** Main sequence relationship between saccade amplitude and duration averaged across all observers. Each point represents one of the nine saccade-amplitude bins with horizontal and vertical error bars indicating ±2SEM. **E** Same as panel D, but each colored line indicating an individual observer’s main sequence fit. **F** lambda waveforms measured at EEG electrode Oz averaged across all participants (see Figure S3 for individual data) for Saccade and Replay sessions (left and right columns, respectively), as well as saccade offset-locked and onset-locked (top and bottom rows, respectively). Colored lines represent average saccade duration in each saccade-amplitude bin (panels D-E). Max-normalized channel topographies (across amplitude bins) for both viewing conditions 90 ms after saccade offset (see Figure S2 for full time course). **G** Response latencies as a linear function of saccade duration in Saccade (dark blue) and Replay (dark red) viewing conditions (upper row: offset-locked; lower row: onset-locked aggregates). Note the strong overlap of the two conditions. **H** Response amplitude as a two-degree polynomial function of saccade duration, same conventions as panel G.

### Identical lambda-response dynamics for real and replayed saccades

After preprocessing and cleaning of recorded data (see Preprocessing and Data cleaning sections), we first ascertained that all observers diligently performed the task in both viewing conditions. Indeed, responses were on average 78.4% (SD = 3.7) correct in the Saccade condition and 77.4% (SD = 4.0) correct in the Replay condition, while no observer exhibited accuracies lower than 70% (Figure 1C). These values were clearly above chance level (*F* (1, 11) = 731.29, 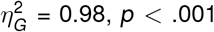) and their differences were not statistically significant (*F* (1, 11) = 1.30, 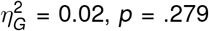), suggesting that any potential differences in task demands had negligible consequences on observers’ task performance.

To investigate to what extent the dynamics of lambda responses depended on saccade metrics, we created nine saccade-amplitude bins, each containing approximately the same number of epochs, for each observer. On average 1332 unique Saccade epochs (range [920, 1561]) entered a bin, representing considerable statistical power that enabled not only robust measurement of lambda responses but also reliable estimation of their latency and amplitude, as we will show below. Note that saccade-amplitude bins, as well as the number of epochs contained in them, were identical in Saccade and Replay sessions due to the direct matching of epochs across viewing conditions (for details, see Data cleaning). Henceforth, unless specified otherwise, the term ‘saccade’ may refer to real and replayed saccades alike. Using nonlinear mixed-effect modeling, we then estimated the relationship between saccade duration and amplitude (Equation 2) which is governed by each observer’s unique saccadic main sequence (Figure 1D-E). Establishing the main sequence at this point will prove an indispensable step for understanding the dynamics of the visual consequences of saccades across varying saccade sizes, as well as for truthful modeling of saccade trajectories.

Figure 1F shows aggregated lambda waveforms for individual saccade-amplitude bins locked to saccade offset (upper row) and saccade onset (lower row) at EEG channel Oz, which displayed the largest response at the time of the strongest global field power, that is, ∼90 ms after saccade offset and ∼125 ms after saccade onset, in both Saccade and Replay conditions (Figure S2). Waveforms clearly show the major feature of lambda responses, that is, a large positive deflection around 100 ms after saccade offset (known as P1 or P100 in EEG) with pronounced occipital topography, suggesting the engagement of early visual areas. Waveforms further exhibit the typical scaling of lambda-response latency with increasing saccade amplitude and duration (Yagi, 1979) and better alignment of waveforms to saccade onset (Amme et al., 2024), but, critically, in Saccade and Replay conditions alike. To determine whether the dynamics of lambda responses were indeed similar in active and passive viewing conditions, we extracted response amplitude (the maximum value of the waveform) and response latency (the time at which 75% of the maximum value is exceeded) from each observer’s individual waveforms (Figure S3) and fitted linear mixed-effects models to predict them based on saccade duration and viewing condition (see Hypothesis tests).

First, we investigated the dynamics of lambda-response latencies. When expressed relative to saccade offset (Figure 1G, top), the mixed model revealed a significant negative overall slope (*β* = −0.61, *t* = −13.62, 95% CI [- 0.71, -0.53]) but no significant difference between viewing conditions, neither in intercept (*β* = 0.22, *t* = 0.20, 95% CI [-2.18, 2.58]) nor slope (*β* = 0.07, *t* = 1.63, 95% CI [-0.01, 0.15], *BF*_10_ = 0.025), suggesting that response latencies decreased with increasing saccade duration in a virtually identical manner in Saccade and Replay conditions. When expressed relative to saccade onset (Figure 1G, bottom), we found the opposite yet slightly weaker pattern: Longer saccades caused an increase of response latency (*β* = 0.36, *t* = 7.64, 95% CI [0.27, 0.45]), further suggesting that lambda responses aligned neither to saccade offset nor to saccade onset. Again, no difference between viewing conditions was found (intercept: *β* = 0.84, *t* = 0.56, 95% CI [-2.01, 3.81]; slope: *β* = 0.05, *t* = 1.14, 95% CI [-0.03, 0.13], *BF*_10_ = 0.014). Importantly, the scaling of response latencies with saccade duration represented a robust finding that held on the level of individual observers (Figure S3): Estimated offset-locked slopes were all negative (range [-0.91, -0.37]) and onset-locked slopes were all positive (range [0.08, 0.58]), suggesting a reliable common source.

Second, we proceeded by characterizing the dynamics of response amplitude. Regardless of whether response amplitudes were extracted from offset- or onset-locked aggregates (Figure 1H, top and bottom, respectively), their relationship with saccade duration was expectedly nonlinear, as evidenced by significant linear terms (offset-locked: *β* = 35.59, *t* = 8.64, 95% CI [27.65, 43.66]; onset-locked: *β* = 48.45, *t* = 12.78, 95% CI [40.56, 56.06]) and quadratic terms (offset-locked: *β* = −20.69, *t* = −5.37, 95% CI [-28.38, -13.23]; onset-locked: *β* = −21.96, *t* = −6.74, 95% CI [-28.54, -15.46]). A significant intercept difference between conditions was present (offset-locked: *β* = 3.86, *t* = 6.58, 95% CI [2.71, 5.03]; onset-locked: *β* = 4.67, *t* = 7.89, 95% CI [3.53, 5.82]), providing clear evidence that response amplitudes were consistently larger in the Saccade than in the Replay condition. Yet, the rate of the increase of response amplitude with saccade duration (linear term of polynomial contrasts) remained largely similar across viewing conditions (offset-locked: *β* = −2.75, *t* = −0.35, 95% CI [-18.11, 12.27]; onset-locked: *β* = 0.38, *t* = 0.05, 95% CI [-15.51, 15.01]), even though its curvature (quadratic term) was less pronounced in the Replay than in the Saccade condition (offset-locked: *β* = −7.13, *t* = −3.45, 95% CI [-11.42, -2.97]; onset-locked: *β* = −11.63, *t* = −5.35, 95% CI [-16.02, -7.18]). This suggests that the effect of saccade size on response amplitude was highly similar but not fully identical in active and passive viewing conditions. Finally, we confirmed the previous finding (cf. Nolte et al., 2025) that response amplitudes were larger if locked to saccade onset than if locked to saccade offset (*M*_*onset*_ = 11.10, *M*_*offset*_ = 10.15, *F* (1, 11) = 41.65, 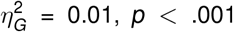), an effect that was more pronounced in the Saccade condition than in the Replay condition (*d*_*Saccade*_ = 1.35, *d*_*Replay*_ = 0.54, *F* (1, 11) = 44.44, 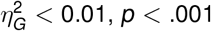).

Some important conclusions can be made at this point. Clearly, lambda responses are not specific to active saccades, but could be elicited by displaying the visual consequences of saccades to the fixating eye. Even though lambda responses following active saccades exhibit larger amplitudes, the functions by which their amplitude and especially latency vary with saccade size were virtually identical in active and passive viewing conditions. This suggests that the systematic variation of response latencies is not, as previously suggested, caused by active predictive processes but by bottom-up processing of the visual change. Finally, lambda responses were locked to neither saccade onset (positive slope) nor offset (negative slope) but to a time point during the saccade – providing a first intuition that their timing is related to the dynamics of saccadic visual consequences.

### Computational modeling of early visual processing explains lambda-response dynamics

To understand whether the dynamics of lambda responses could be understood purely in terms of early visual processing, we set up a large-scale simulation to model saccadic visual shifts. To that end, we first approximated observers’ visual surroundings, that is, the natural scene presented on an otherwise empty monitor with a dark uniform background (Figure 2A). We simulated six horizontal saccade trajectories (1.5–14 dva amplitude) matching observers’ average main sequence, as well as random ocular drift movements (for details, see Computational modeling), and aligned them in a way that start and end points of saccades were balanced (Figure 2B). According to these trajectories, we simulated retinal input over time, which served as model input (Figure 2C). Importantly, retinal changes included either movement of both scene and screen as gaze traversed the entire visual field (Saccade condition), or movement of only the scene relative to the screen as gaze remained fixed (Replay condition). The model (Figure 2D) featured first a spatial filtering stage where retinal images were convolved with a bank of Gabor filters, then a temporal filtering stage where resulting filter responses were constrained to the temporal sensitivity profile of the human visual system (Kelly, 1979), and finally, after squaring of quadrature-paired filter responses, a delayed-normalization phase that gave rise to transient-sustained response dynamics representative of the visual cortex (Groen et al., 2022). To compute evoked response-like time series, as shown in Figure 2E, we summed model responses across spatial-frequency (SF) and orientation channels, as well as across space. Note that the model was not fitted on any of our experimental data. The model’s parameters were unchanged from what was initially derived from neurophysiological and psychophysical evidence (Schweitzer, Seel, et al., 2025) and simulations were performed with stimuli shown during the experiment using saccade metrics similar to those found in the data (Figure 1D). Strikingly, lambda responses, with peaks occurring around 114 ms (SD = 14.5) after saccade onset, emerged reliably from the model’s output (Figure 2E).

**Figure 2.**
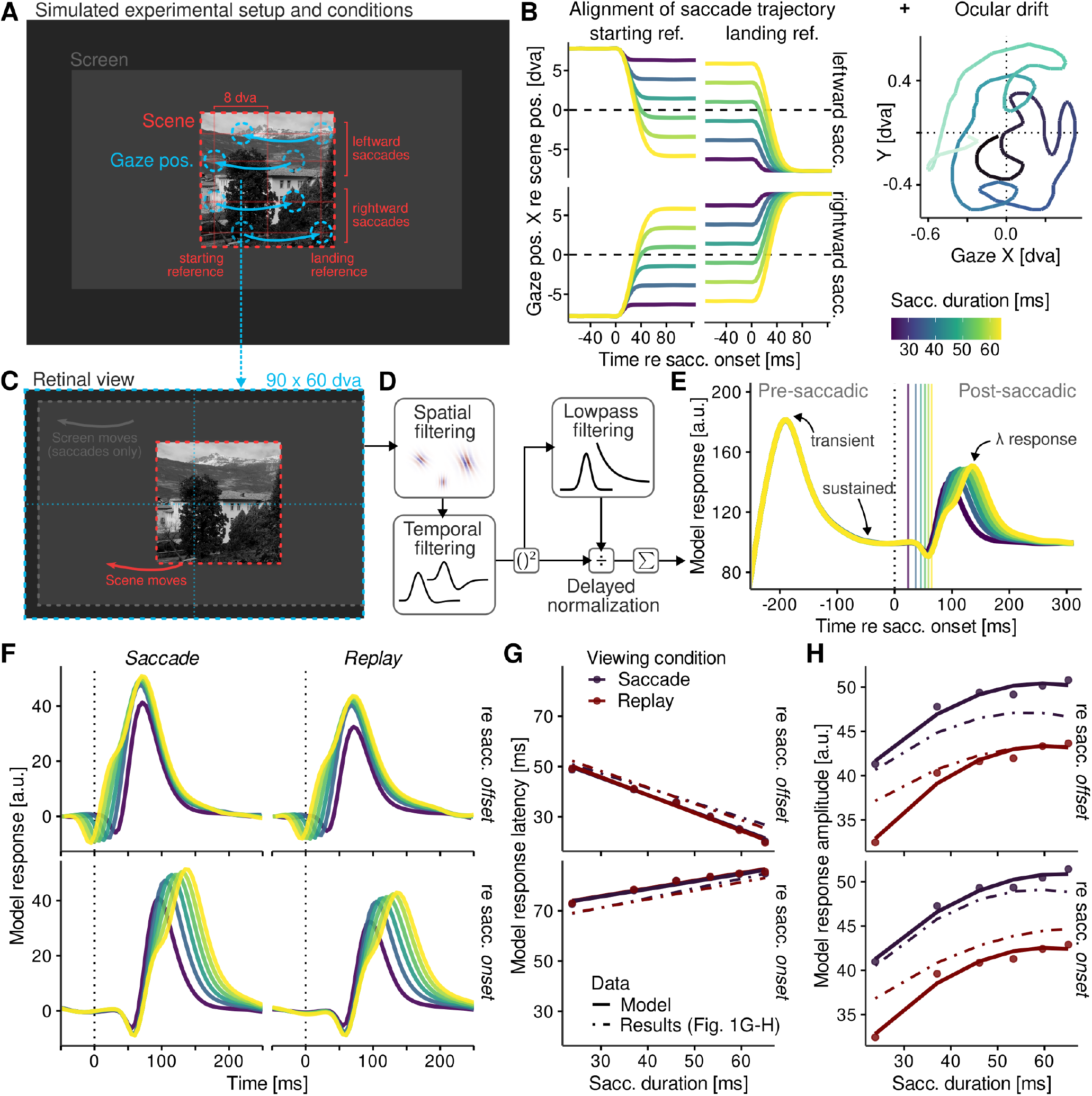
Modeling of early visual responses to real and replayed saccades. **A** Simulated experimental screen (grey filled rectangle) with scene (red dashed outline) on uniformly black background. Simulated gaze trajectories (blue) are illustrated in four rows for leftward vs rightward saccade directions and starting vs landing references, respectively (see Computational modeling for details). **B** Simulated saccade trajectories (left) and ocular drift trajectory (right) used for the simulation. **C** Example one retinal view of an entire temporal sequence of views which was used as model input. **D** Basic model processing steps (for details, see main text and Schweitzer, Seel, et al., 2025). **E** Prototypical model output (averaged across conditions shown in panel B) for six saccade amplitudes. Note that both the transient and sustained phase of the pre-saccadic response (i.e., to the onset of the simulated screen) was identical for each amplitude. Solid vertical lines denote saccade offsets. **F** Simulated lambda responses, baseline-corrected and averaged across all tested stimuli. Plotting conventions same as Figure 1F. **G-H** Response latencies and amplitudes as functions of simulated saccade duration (solid lines). Dot-dashed lines in each panel show fits of experimental results presented in Figure 1G-H, but with mixed-model grand intercepts adjusted by 32.7 ms to account for differences in absolute latencies (offset-locked: β_0_ = 33.42 compared to β_0_ = 69.64 measured; onset-locked: β_0_ = 81.08 compared to β_0_ = 110.22 measured), as temporal response functions in the model did not include any explicit delays (cf. Reich, Mechler, & Victor, 2001, their Fig. 2B).

If latency and amplitude of lambda responses depended on the dynamics of the saccade-induced visual change, then model responses should change in a way similar to measured responses. Indeed, average model-response waveforms (Figure 2F) were highly similar in Saccade and Replay conditions and appeared to vary systematically with saccade duration. As with measured responses, we extracted model response latency and amplitude from these waveforms and fitted linear regressions. First, we found that the slopes describing changes of response latency were remarkably similar to previous measurements (Figure 2G), both for offset-locked (*β* = −0.69, *t* = −31.8, *p <* .001; cf. *β* = −0.62 with SE = 0.045 measured) and onset-locked aggregates (*β* = 0.30, *t* = 13.0, *p <* .001; cf. *β* = 0.36 with SE = 0.047 measured), and remained unaltered across viewing conditions (offset-locked: *β* = 0.02, *t* = 0.57, *p* = .587; onset-locked: *β* = −0.01, *t* = −0.11, *p* = .915). Overall, fits of model predictions and measurements were nearly perfectly correlated (offset-locked: *r* (10) = 0.99, *p <* .001; onset-locked: *r* (10) = 0.99, *p <* .001). Second, we also found pronounced similarities in between modeled and measured response amplitudes (Figure 2H). Model-response amplitudes increased nonlinearly with saccade duration, as both significant linear (offset-locked: *β* = 15.53, *t* = 11.82, *p <* .001; onset-locked: *β* = 15.82, *t* = 13.4, *p <* .001) and quadratic terms (offset-locked: *β* = −6.21, *t* = −4.73, *p* = .003; onset-locked: *β* = −5.55, *t* = −4.70, *p* = .003) of polynomial contrasts were significant. The model also replicated larger response amplitudes in the Saccade condition than in the Replay condition (offset-locked: *β* = 7.54, *t* = 14.07, *p <* .001; onset-locked: *β* = 8.21, *t* = 17.03, *p <* .001), while both linear (offset-locked: *β* = −2.84, *t* = −1.08, *p* = .320; onset-locked: *β* = 0.01, *t* = 0.01, *p* = .996) and quadratic terms (offset-locked: *β* = 0.91, *t* = 0.35, *p* = .740; onset-locked: *β* = 0.92, *t* = 0.39, *p* = .709) remained invariant across viewing conditions. Despite unit-related differences in intercept, fits of model predictions and measurements were again highly correlated (offset-locked: *r* (10) = 0.98, *p <* .001; onset-locked: *r* (10) = 0.97, *p <* .001), suggesting that amplitudes of lambda responses, too, can be approximated well enough in terms of visual-only mechanisms.

Even though its output emerged solely from a passively receiving, feedforward type of visual processing, the model succeeded at closely reproducing latency-related and amplitude-related dynamics. Notably, the prominent differences in lambda-response amplitude between viewing conditions were likely caused by visual differences: In the Saccade condition saccade-induced transients result from combined motion of the scene, the screen and its surroundings, whereas in the Replay condition the scene alone moves while screen and surroundings remain static. The movement of screen edges in the Saccade condition stimulates peripheral receptive fields, especially those selective to low SFs which are known to only reach their optimal temporal-frequency (TF) range at very high velocities (Burr & Ross, 1982). Moreover, sharp screen edges, with their step-like spatial profile, elicit responses in all spatial channels and should therefore produce larger overall responses. We compared scene stimuli with sharp and soft edges (for details, see Stimuli) and found that sharp edges indeed produced stronger onset-locked model responses (*M*_*sharp*_ = 44.01, *M*_*soft*_ = 39.67, *t* (11) = 10.53, *p <* .001). This effect could be confirmed, albeit much weaker, in EEG measurements (*M*_*sharp*_ = 11.14 *µV*, *M*_*soft*_ = 11.07 *µV*, *F* (1, 11) = 5.17, 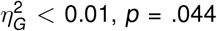), suggesting that low-level features of visual stimuli may well influence lambda responses as their generation is fundamentally grounded in early visual processing.

### The natural visual consequences of saccades give rise to lambda responses

Both experimental measurements and computational modeling we have revealed remarkably similar latency dynamics in active and passive viewing conditions. To explain these dynamics, three crucial elements are required. First, we computed the velocity profiles of the simulated saccade trajectories used previously (Figure 3A). According to the observers’ average main sequence, saccadic duration and peak velocity (both closely related; Figure S4A) increase systematically with saccade amplitude. Second, we used the psychophysically measured spatiotemporal contrast sensitivity surface estimated by Kelly (1979) to approximate how SF-specific contrast sensitivity changes depending on the TF or velocity at which the image moves (Figure 3B). Third, we extracted the average slope of the ^1^*/*_*f*_ power law by which log radial-average spectral amplitude decreased with increasing log SF in our scene stimuli (*β*_*SF*_ = −1.36, SE = 0.02; Figure 3C). This estimated slope was highly similar to previous estimates from natural scenes (i.e., between -0.8 and -1.5; Tolhurst, Tadmor, & Chao, 1992).

**Figure 3.**
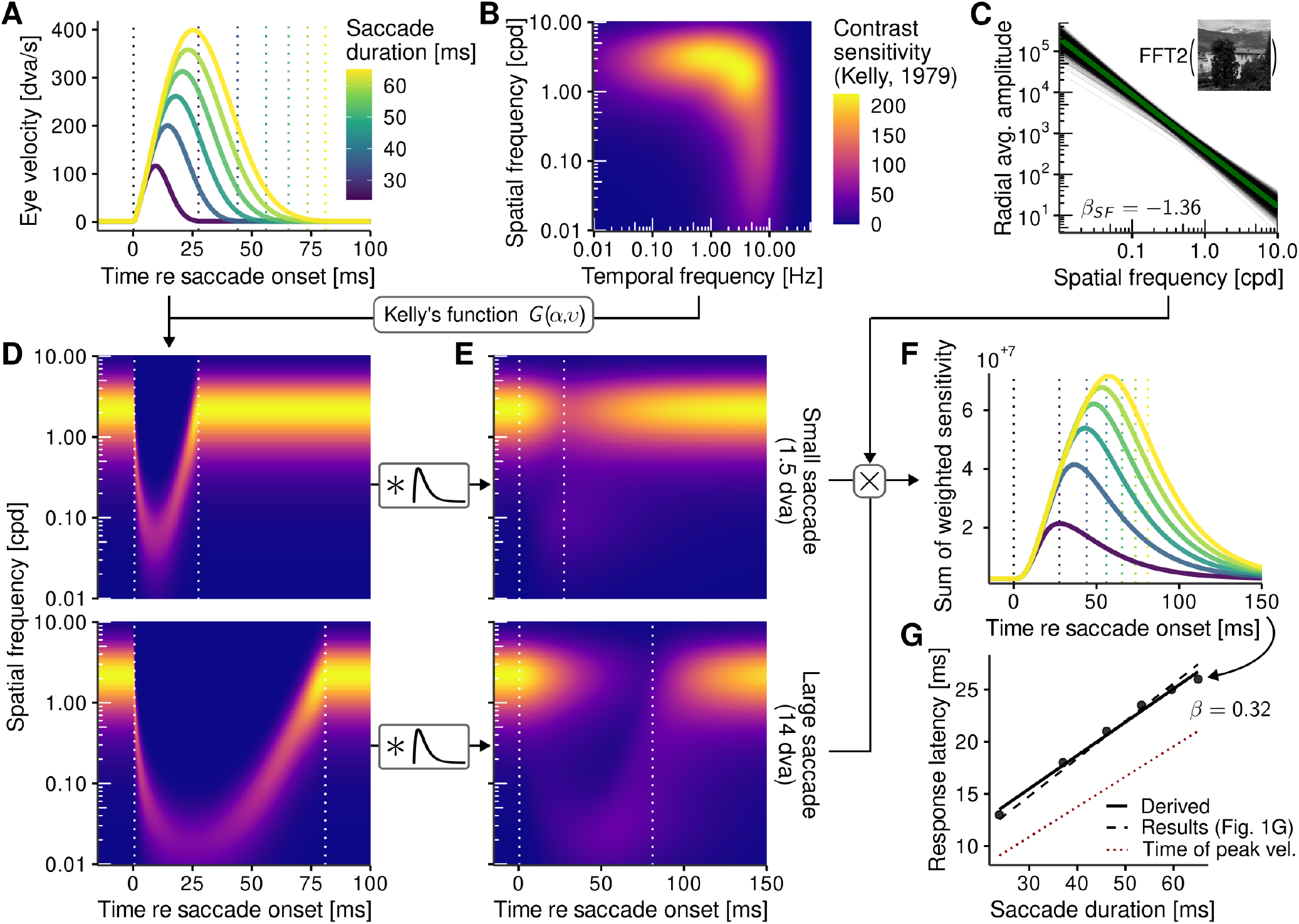
Mechanistic explanation for the latency scaling of lambda responses. **A** Velocity profiles of six saccades simulated by means of Equation 5. Dotted vertical lines indicate the offsets of these saccades when detected based on values falling below a velocity threshold of 1.3 dva/s. **B** Spatiotemporal sensitivity surface described by Kelly (1979), computed by G(α, υ) where α = 2π · spatial frequency and υ refers to retinal velocity (see Equation 4). **C** 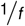 power-frequency relationship determined by fitting a mixed-effects log-linear model on stimulus scenes’ radial amplitude spectra. Thin black lines indicate estimates for individual scenes, the thick green line their average. **D** Saccade-induced change of contrast sensitivity over time, shown for the smallest (1.5 dva) and largest (14 dva) visual change. **E** Same as panel D, but after convolution with a gamma-shaped temporal response function (see Computational modeling). **F** Product of convolved contrast sensitivity and each corresponding spatial frequency’s average amplitude estimate (panel C) summed across spatial frequencies [a.u.]. Conventions same as panel A. **G** Response latency, extracted from summed contrast sensitivity (panel F), as a function of simulated saccade duration (solid line). The black dashed line indicates the intercept-corrected slope reported in Figure 1G, bottom row, collapsed across viewing conditions, whereas the red dotted line indicates the time of the saccadic peak velocity averaged across observers (for individual estimates, see Figure S4).

Having established these elements, we computed theoretical contrast sensitivity profiles over time for all velocity profiles from small to large saccades (Figure 3D). Two outcomes should be noted: First, as the velocity of the retinal shift increases, the sensitivity profile at fixation breaks down and gradually shifts to lower SF bands. This outcome was also observed across the various spatial-frequency channels employed in our computational model (Figure S5, see also SF-specific model-response dynamics in the Supplementary Material). Second, as a consequence of the main-sequence relationship, larger (and thus faster) saccades reach lower SF bands and maintain sensitivity in these bands for a longer time. By further convolving these time-resolved sensitivity profiles with a suitable temporal response function we simulated temporal integration, creating a more realistic representation of velocity-dependent changes in contrast sensitivity (Figure 3E). As a final step, sensitivity at each SF and time point was weighted by the scenes’ estimated power spectrum and summed across SFs, giving rise to theoretical response profiles (Figure 3F). The latencies extracted from these profiles increased linearly with saccade duration in nearly the same way as measured lambda-response latencies (*β* = 0.32, *t* = 19.37, *p <* .001, cf. *β* = 0.36 measured; Figure 3G), suggesting that the three elements outlined above were in principle sufficient to explain the observed latency dynamics of lambda responses.

This proposed explanation makes two testable predictions. First, if latency dynamics are dominated by saccade-induced visual transients in the low-SF domain, then latencies should be best explained by the time when these transients are strongest, that is, when the saccade’s velocity reaches its peak. For each single observer, we thus determined (relative to saccade onset) the time of saccadic peak velocity, revealing that it was closely related to the saccade’s duration (*β* = 0.29, *t* = 11.00, 95% CI [0.24, 0.34]; Figure S4B). Remarkably, this slope was also highly similar to the slope describing the latencies of lambda responses (cf. red dotted line in Figure 3G). Furthermore, observers’ individual lambda-response latency estimates were predicted extremely well by their individual times of peak velocity (*β* = 1.32, *t* = 6.72, 95% CI [0.92, 1.69]), regardless of viewing condition (*β* = 0.16, *t* = 1.06, 95% CI [-0.12, 0.48]). With respect to goodness of fit, time of peak velocity and saccade duration were equally good predictors (BIC_t(vpeak)_ = 993.65, BIC_dur_ = 993.78). Second, as higher saccadic peak velocities should produce stronger transients in the low-SF domain, one would expect that onset-locked lambda-response amplitude would be best explained by the peak velocity value itself. This was exactly the case (BIC_vpeak_ = 696.7, BIC_dur_ = 704.1, *BF*_10_ = 40.4), corroborating the functional relevance of saccade dynamics for driving lambda responses.

Finally, to test whether lambda responses are statistically locked to the time of peak velocity, we corrected observers’ individual latency estimates by subtracting their corresponding times of peak velocity. Following this correction, the overall slope was reduced to 0.07 (range [-0.21, 0.34]) which was not significantly different from zero (*t* = 1.31, 95% CI [-0.04, 0.17]), regardless of viewing condition (*β* = 0.05, *t* = 1.16, 95% CI [-0.03, 0.13]). Model comparisons strongly favored the intercept-only model (BIC_null_ = 985.71, BIC_dur_ = 997.29, *BF*_01_ = 327.0), suggesting that corrected lambda-response latency no longer scaled with saccade size. In other words, lambda responses best locked to the time of saccadic peak velocity where the strongest saccade-induced visual transients were expected to arise. These results not only corroborate the proposed mechanistic explanation for lambda responses, but also suggest a unified, theoretically grounded proxy for their latency, that is, 96.3 ms (*t* = 69.1, 95% CI [93.6, 98.9]) after the time of saccadic peak velocity.

## Discussion

More than 70 years after their discovery (Evans, 1953; Gastaut, 1951), here we provide a model-based explanation of how lambda responses are generated. We collected several thousands of saccade epochs of EEG data during active free-viewing of natural images and compared them to an experimental condition where those saccades’ visual consequences were replayed during fixation. We did not only find virtually similar lambda-response dynamics in the two conditions, but the same response dynamics emerged from a purely early-visual computational model. Using this modeling as a catalyst, we extracted the basic principles needed for the observed lambda-response dynamics to arise and confirmed their empirical predictions, establishing the new hypothesis that lambda responses are most parsimoniously explained by the visual dynamics that naturally occur as consequences of saccades.

First, the dynamics of lambda responses were virtually identical in Saccade and Replay conditions, regardless of whether waveforms were computed from saccade offset-locked or onset-locked aggregates. This finding demonstrates that the observed scaling of lambda-response latency was in fact not a result of extra-retinal mechanisms involved in active vision, but must be linked to what is common among conditions: Bottom-up, feedforward visual processing. It should be noted that, despite our best efforts to keep the retinal input as comparable as possible, there is no perfect replay condition, because it is impossible to reproduce either the movement of the presentation monitor and visual references around it, the exact dynamics of eye movements as measured by video-based ET, or the forces that saccades exert on the lens (Deubel & Bridgeman, 1995) and the retina (Richards, 1969). Clearly, we were also unable to perform retinal stabilization during Replay sessions. Despite that, latency dynamics in the Replay, and to a lesser degree also amplitude dynamics, were statistically indistinguishable from the Saccade condition. That simulated saccades can produce responses similar to saccadic lambda responses has been shown in human EEG (Barlow & Cigánek, 1969; Lesèvre & Remond, 1972), as well as in the optic tract, lateral geniculate nucleus, and visual cortex of the cat (Ebersole & Galambos, 1973), but a fully matched comparison of real and simulated saccades in a free-viewing setting using natural stimuli has, to our knowledge, never been attempted. With our focus on the visual modality, which clearly matched the topography of the EEG signal (cf. Figure S2), we have purposefully not looked at any other potential differences between the two viewing conditions. In principle, our results do not speak against the involvement of extra-retinal mechanisms. On the contrary, it is obvious that the mere phenomenological difference between Saccade and Replay conditions – in the Saccade condition the scene is static, in the Replay condition it moves – is decisive evidence that the visual system must take oculomotor information into account. However, this difference does not seem to be captured in the first feedforward sweep through the visual cortex, i.e. in the lambda response, but likely arises from a later stage of perceptual processing. This later stage might already start around 150 ms, further downstream in the visual pathway (Huber-Huber, Buonocore, Dimigen, Hickey, & Melcher, 2019) where neural activity might be maintained across individual fixations to provide trans-saccadic predictions (Huber-Huber, Buonocore, & Melcher, 2021; Parr, Sajid, Da Costa, Mirza, & Friston, 2021; Valsecchi & Gegenfurtner, 2016).

Second, measured dynamics of lambda responses, including similarities and differences between viewing conditions, could almost completely be reproduced by a computational model that included basic mechanisms of early visual processing. The model was neither built for specifically predicting lambda responses (Schweitzer, Seel, et al., 2025) nor fitted in any way to the data. Neither model nor simulations included every potentially important aspect of the human visual and oculomotor system. For instance, the model did not include the SF-specific reduction of sensitivity at increasing eccentricities (e.g., Pointer & Hess, 1989) or contrast-dependent latencies of V1 simple cells (Reich et al., 2001). Moreover, simulations did not include visual structure around the presentation screen. Simulated saccade trajectories also did not include postsaccadic oscillations that are prominent in video-based ET (Schweitzer & Rolfs, 2022) and have been shown to reliably affect visual localization judgments (Deubel & Bridgeman, 1995), as well as the perception of smear induced by replayed saccades (Schweitzer, Doering, Seel, Raisch, & Rolfs, 2025). The model only needed human observers’ saccade dynamics, their stimulus display, and especially the psychophysically estimated spatiotemporal sensitivity profile of the human visual system (Kelly, 1979), to produce the phenomenon of latency scaling of the lambda response. The computational model also qualitatively reproduced amplitude differences between Saccade and Replay conditions, suggesting that it would be premature to attribute this difference solely to the difference in active and passive viewing.

Third, and finally, we proposed a parsimonious and visual-only explanation for the latency dynamics of lambda responses that approximated the measured latencies very closely. The core idea is that saccades introduce considerable velocities to the retinal image, thereby greatly modulating visual spatiotemporal sensitivity. On the one hand, high SFs are rendered invisible as saccadic velocities introduce TFs unresolvable by the visual system (Castet, Jeanjean, & Masson, 2002; Schweitzer, Doering, et al., 2025). On the other hand, low SFs are boosted as saccadic velocities shift them into their optimal TF range (Burr & Ross, 1982). It is fair to say that saccade-induced retinal shifts represent the only instance when such low-SF transients are introduced during normal vision, which is why they are considered highly important for the visual extraction of low-SF information (Boi, Poletti, Victor, & Rucci, 2017). The larger the saccade, the higher the power of luminance modulation in the low-SF domain (Mostofi et al., 2020), and preliminary evidence confirms that larger saccades also lead to higher contrast sensitivity to low-SF stimuli, notably in both real and simulated saccades (Howard Li, Cox, Victor, & Rucci, 2025). For viewing natural scenes, whose spectral content is subject to the ubiquitous ^1^*/*_*f*_ power law (Tolhurst et al., 1992), this is critical: Larger (and thus faster) saccades are capable of boosting lower SFs which have higher power in the visual image, thus producing greater overall transients (cf. Figure 3F). This mechanism, which is a mere consequence of the spatiotemporal sensitivity profile of the human visual system (Kelly, 1979), explains the increase and saturation of lambda-response amplitudes with saccade size: Transients increase as saccadic velocities gradually enhance low-SF bands, but also saturate as the saccadic main sequence saturates. Importantly, the strength of the saccade-induced visual transient should not only vary between saccade sizes but also as a function of any given saccade’s velocity profile, that is, it should be strongest around the time of the peak velocity. In other words, as the latency of the peak velocity scales with saccade size due to the main sequence, so should the latency of the lambda response. This was precisely what emerged from our computational modeling as well as from the EEG data. While this suggests that lambda responses are dominated by low-SF transients, it also leaves an important role for saccade offsets: Once the eye is fixating, the typical contrast sensitivity profile is reestablished and gives rise to offset-locked visual transients in the high-SF domain. Interestingly, this saccade-offset locked component was also predicted by the model (Figure 2F) and might correspond to the later and much smaller N1 component of the lambda wave (at ∼170 ms) which has been shown to be more closely time-locked to the saccade offset (Yagi, 1979) and could therefore reflect the later stages of the coarse-to-fine progression of visual processing that naturally emerges from the oculomotor cycle (Boi et al., 2017).

We found that visual consequences of saccades explain the dynamics of lambda responses – regardless of whether actual saccades were executed – given three ingredients: the human spatiotemporal visual sensitivity profile, the main sequence as a global law of oculomotor control, and natural scene statistics. This finding speaks to the long-standing question of how much and what types of the visual input reaching the retina during saccades is actually processed by the brain (Binda & Morrone, 2018; Matin, 1974; Ross, Morrone, Goldberg, & Burr, 2001; Volkmann, 1986). During natural viewing we have little to no perceptual awareness of the visual consequences of our own saccades, that is, we perceive neither saccade-induced motion and smear (saccadic omission; Campbell & Wurtz, 1978) nor irregular jumps or displacements of the world as eye movements shift the visual image across the retina (visual stability; Bridgeman, Van der Heijden, & Velichkovsky, 1994). It might thus appear surprising that visual input during saccades rather than from new fixation trigger the first and most prominent saccade-locked EEG response. This notion, however, does not have to be at odds with the phenomenal differences between active and passive viewing conditions: In the Saccade condition, motor signals about imminent eye movements allow for an anticipatory remapping of visual space, whereas during Replay retinal changes remain entirely unpredictable – highlighting the critical role of efference-copy signals in achieving visual stability (Wurtz, 2018). Although these mechanisms eventually must affect the visual phenomenology, they do not appear to alter the early visual processing of the saccadic shift – a notion supported by neurophysiological data recorded from monkey V1 (Ilg & Thier, 1996), as well as recent behavioral evidence from the perception of motion smear (Schweitzer, Doering, et al., 2025).

To conclude, after experiments conducted several decades ago have already converged on the conclusion that lambda responses are fundamentally visual potentials, understanding lambda responses in terms of saccade-induced visual transients not only enables the prediction of their specific dynamics, but also explains why they were found to be largely similar in real and simulated saccades. Given their reliable measurement, as well as tractability via computational modeling, lambda responses may prove a valuable tool for understanding visual processing around the time of eye movements. Conversely, endowing the present computational model with justified assumptions about the generation of neurophysiological signals in the brain may pave the way towards predicting actual visual evoked responses measured with EEG, MEG, or intracranial techniques.

## Methods

### Participants

Twelve observers (9 female, mean age 25.3, age range [19, 35], 10 right-handed, 4 right ocular dominance), of which one was also an author of this study, participated in the experiment. All gave written informed consent and had normal or corrected-to-normal vision. The experiment was pre-registered (https://osf.io/jn7z6), conducted in agreement the Declaration of Helsinki (2013), and approved by the Research Ethics Committee at the University of Trento, Italy (protocol number 2023-034).

### Stimuli

72000 quadratic color images were extracted from the Natural Scene Dataset (Allen et al., 2022) along with their object annotations contained in the COCO database (Lin et al., 2014). Based on these annotations, we selected only those images without semantic loss after image cropping and with more than five different annotations of different object categories. Images with artificial frames or borders were manually flagged and removed, resulting in a final selection of 1980 images and annotations, made publicly available at https://osf.io/3txsd. Selected images were upsampled to 650^2^ pixels using bilinear interpolation and padded with the image pixels’ average RGB value, resulting in a image size of 725^2^ pixels (approx. 21.3 dva). To create soft image borders a two-step procedure was applied: First, a copy of the image was convoluted with a Gaussian kernel (*σ* = 7.5 pixels) and, second, the original image was mixed with its blurred version using an alpha mask created by convolution of the image outline with the same Gaussian kernel. White circles with 0.2dva radius and 1-pixel line width served as fixation dots. During replay, when fixation dots remained visible, their alpha was reduced from 1.0 to 0.5 in order to reduce their saliency.

### Apparatus

The experiment was carried out in a sound-attenuated and electromagnetically shielded cabin. Stimuli were shown on an 26.5-in ASUS ROG Swift PG27AQDP OLED monitor (gamma = 2.2, maximum brightness = 400 cd/m^2^) with a G-Sync-enabled variable refresh rate of up to 480 frames per second and a spatial resolution of 1920 x 1080 pixels, connected to a NVIDIA GeForce RTX 5070 graphics card. To ensure the fidelity of stimulus presentations, we systematically validated the timing of our setup (see Evaluation of stimulus timing in the Supplementary Materials for details). Observers’ heads were supported by a chin rest at a distance of 59.5 cm from the monitor. Stimulus display was controlled using Psychtoolbox 3.0.19 (Kleiner et al., 2007) on Matlab 2025b (Mathworks, Natick, MA, USA) in Ubuntu 24.04. Monocular eye tracking of the dominant eye was performed with an Eyelink 1000 desktop system (35-mm lens, leveled mode, link heuristic-filter level 2) controlled via the Eyelink Toolbox (Cornelisen, Peters, & Palmer, 2002). 64-channel EEG data was collected with a BrainAMP DC system (Brain Products, Gilching, Germany) at a sampling rate of 1000 Hz and applying 250-Hz low-pass and 0.016-Hz high-pass recording filters. We had a standard actiCAP layout available, but used electrodes FT9 and FT10 as left and right horizontal EOG (outer canthi), as well as electrodes T7 and T8 as left and right vertical EOG (infraorbital), respectively (https://osf.io/crh8y/files/62w79). Triggers of 2 ms duration were concurrently sent to Eyelink host and EEG system, allowing the temporal alignment of ET and EEG data using the EYE-EEG Toolbox (Dimigen et al., 2011) in EEGLAB (Delorme & Makeig, 2004). Responses were given with a standard US-english USB keyboard.

### Procedure

The experiment consisted of four separate sessions, that is, two Saccade sessions and two subsequent Replay sessions. Naive observers first completed both Saccade sessions before they were introduced to the Replay task in their third session. There were nine blocks of 110 trials/images per session, of which each was preceded by a nine-point calibration of the eye tracker. Each block contained 55 images with sharp edges and 55 images with soft edges (see Stimuli for details), presented in random interleaved order. This order was re-randomized within each block of the Replay sessions. Which images were assigned to which edge condition was counterbalanced across observers. Scene images presented during Saccade sessions were reused in Replay sessions and were presented in the same edge condition. On average, 9 days (range [1, 18]) passed between the first Saccade and first Replay session and 14 days (range [1, 33]) passed between the second Saccade and second Replay session. By design, at least 1870 trials/images were administered before any scene was repeated during Replay.

#### Saccade sessions

Trials started with a fixation control requiring observers to fixate within a circular area (*r* = 1.25 dva) around a single fixation dot for 350–450 ms, allowing 80 breaks in fixation and 2 seconds without valid fixation before triggering a calibration and rescheduling the trial at the end of the block. Locations of fixation dots were uniformly sampled from a 17^2^-dva grid centered around screen center to systematically vary from where observers started exploring the visual scene. Upon successful fixation the fixation dot was extinguished and the scene image was presented at screen center for 2.5 seconds. After a blank interval of 100 ms, a response cue, consisting of the name of an object category and the two response options (see example in Figure 1A), was presented with a 1/8 probability. In this case, observers responded whether the object was present (RightArrow key) or absent (LeftArrow key). Ground-truth object presence was based on object annotations contained in the COCO database (see Stimuli) and was randomly sampled with equal probability.

#### Replay sessions

Fixation control was identical to the Saccade session with the exception that fixation dots were sampled from a much smaller 3^2^-dva grid around screen center. Once fixation control was passed, the presentation of the replay sequence (see Replay trajectories for details) was initiated, causing the scene image to rapidly shift across the scene according to previously recorded eye movements. In order to help observers to hold fixation during presentation, the fixation dot remained on the screen with reduced opacity. After a blank interval of 100 ms, observers were prompted to perform the same task as during Saccade sessions but on a different, randomly sampled subset of trials.

#### Replay trajectories

Replay trajectories were based on gaze positions retrieved online during Saccade sessions at a (not necessarily uniform) sampling rate of 1000 Hz. To greatly reduce the impact of ET noise on replay trajectories – which would become disturbingly evident as a high-frequency jitter of the visual image – we developed a dedicated pipeline to preprocess gaze-position data. As a first step, the gaze-position signal was padded with 1000 samples on each side and resampled to 480 Hz. Bandlimited downsampling was used to not only removed high-frequency noise from the signal but also create uniformly sampled data, critical for the next step. Second, velocity-based saccade detection (Engbert & Kliegl, 2003, *λ* = 6, minimum duration: 6 samples) was performed. Resulting saccade candidates were merged if they were separated by less than 10 samples, thus combining the main saccade with the post-saccadic oscillation that is likely to occur due to the inertial movement of the iris (Schweitzer & Rolfs, 2022). Third, two moving-median filters, that is, the fixation filter with a span of 51 samples and the saccade filter with a span of only 5 samples, were applied to gaze-position data. The two resulting filtered signals were then averaged using an event vector (0: fixation; 1: saccade) specifying weights over time. In this event vector saccade segments were expanded by five samples on each side before convolution with a Gaussian kernel (span: 10 samples) achieved a smooth and continuous transition between fixation and saccade segments. This procedure achieved efficient noise reduction during fixation periods while completely avoiding distortions of saccade trajectories. Fourth, a drift correction was performed on the filtered signal by subtracting the offset between fixation dot and initial gaze position (i.e., median gaze position prior to passing fixation control). Finally, the filtered, drift-corrected signal was cropped to the relevant stimulus-presentation interval while taking into account our display’s video delay (Figure S1C), resulting in the pre-processed gaze-position signal *G*_*x*,*y*_ used during Replay.

During Replay, scene position in screen coordinates at a given time point *R*_*x*,*y*_ (*t*) was computed according to

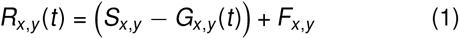

where *S*_*x*,*y*_ is the (constant) scene position during Saccade sessions and *F*_*x*,*y*_ is the instructed fixation position during Replay sessions. While the latter was by default constant, it could be updated to the observer’s current gaze position, that is, the median of the 15 most recent gaze-position samples, if gaze was found outside of a circular fixation boundary (*r* = 2.5 dva) around the instructed fixation position. This gaze contingency not only served as a disincentive to making catch-up saccades, but also allowed replay dynamics to remain comparable across varying fixation positions. Furthermore, *G*_*x*,*y*_ (*t*) was linearly interpolated from the pre-processed gaze-position signal, supplying *t* as the current time passed since presentation onset. By means of this approach durations of replay sequences were kept equal to presentation durations in their corresponding Saccade trials, efficiently taking into account variable frame durations and refresh rates slightly lower than 480 Hz (see Figure 1B and Evaluation of stimulus timing).

### Analysis

#### Preprocessing

EEG data was first highpass-filtered (cutoff: 0.025 Hz) and an automatic channel rejection (Bigdely-Shamlo, Mullen, Kothe, Su, & Robbins, 2015, flat line criterion: 2 s, line noise criterion: 7 standard deviations, minimum channel correlation: 0.4) was performed. On average 1.3 (range [1, 4]) bad channels were detected in 11.8% of all blocks. We furthermore screened for bridged channels using the eBridge algorithm (Alschuler, Tenke, Bruder, & Kayser, 2014, with default parameters) and detected one pair of bridged channels in all sessions. Spherical interpolation was used to reconstruct bad and bridged channels. Finally, data was then lowpass-filtered (cutoff: 98 Hz) and line noise (50 Hz) was removed using the EEGLAB plugin CleanLine (Mullen, 2012) with a bandwidth of 2 Hz, upon which an average reference (including EOG electrodes) was computed.

EEG and ET data, both recorded with a sampling rate of 1000 Hz, was aligned using the EYE-EEG toolbox (Dimigen et al., 2011) and saccades were detected using the Engbert-Kliegl algorithm (Engbert & Kliegl, 2003, *λ* = 6, minimum duration: 12 samples, merge interval: 21 samples). To remove artifacts related to eye movements, an optimized independent component analysis (ICA) procedure (Dimigen, 2020) was applied. As recordings were stopped between blocks to allow for potential adjustments of electrode impedance, one ICA per block was trained on continuous data, that is, cropped from the first calibration of the eye tracker to the end of the last trial in each block. The data underwent highpass filtering at a 2-Hz cutoff and an overweighting (100%) of spike potentials retrieved from the data around saccade onset ([−20, +10] ms). ICA weights were then applied to the original EEG data, finally allowing for an automatic rejection of ocular ICs based on a saccade/fixation variance ratio threshold of 1.1 (Plöchl, Ossandón, & König, 2012). No manual rejection of ICs was performed beyond that.

Replay events – that is, onset and offset timestamps of replayed saccades, as well as scene position and eye positions over time – were added to the combined EEG and ET data. This step was critical to achieve a one-toone matching between real-saccade epochs in the Saccade sessions and their corresponding simulated-saccade epochs in the later Replay sessions. Validating this matching procedure we found that saccade-onset detection performed during the creation of replay trajectories (see Replay trajectories) produced event timestamps highly similar to saccade onsets detected offline during preprocessing (median distance: 3 ms, median absolute deviation: 1 ms). This suggests not only that replay trajectories were created with high temporal precision based on online-sampled ET data during Saccade sessions, but also that post-hoc synchronization of replay trajectories with recorded EEG-ET data was accurate. During preprocessing, entire trials were automatically excluded either if no saccades could be detected or if participants erroneously responded without being prompted, thereby preemptively ending the trial. These rare cases only ever occurred in four out of 432 blocks. Detected saccades were excluded if their ET data contained any missing values. Finally, epochs relative to either real or simulated saccade onset and offset ([-50, 300] ms, with [-50, 0] used for baseline correction) were created from combined EEG and ET data. To be included, saccades had to be initiated more than 50 ms prior to the end of the scene’s presentation. In order to reduce working-memory load, epochs were padded and downsampled to 200 Hz using bandlimited interpolation.

#### Data cleaning

Cleaning was performed in six sequential exclusion steps. First, entire replay trials were removed in case of erroneous drift correction during the generation of replay trajectories caused by blinks during fixation control in Saccade sessions – a bug that was fixed during data collection and therefore only affected four observers and 1.9% (range [0.1, 6.2]) of their trials. Second, 1.5% (range [0.2, 3.2]) epochs in Saccade sessions and 0.5% (range [0.1, 1.7]) in Replay sessions were removed if missing ET samples, most prominently due to blinks, were contained in the data. Third, Replay epochs were excluded if saccades or blinks were made around the onset of replayed saccades, that is, [-50, 200] ms. This case occurred more frequently (*M* = 8.6%, range [3.6, 29.8]) due to automatic catch-up saccades in response to rapid displacements of the scene. Fourth, epochs with unrealistic saccade metrics were excluded. To detect such metrics, individual main sequence functions were fitted using the two-parameter exponential function

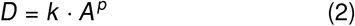

where *D* is saccade duration and *A* is saccade amplitude (Becker, 1989). If absolute residual values were larger than three standard deviations of the residual distribution or saccade amplitudes larger than 20 dva, epochs were rejected, amounting to 1.1% (range [0.1, 2.0]) of the data. Fifth, to make sure both viewing conditions were balanced in terms of underlying saccade metrics and set size, only those epochs were kept for which both Saccade and Replay epochs remained valid after the previous exclusion steps (*M* = 92.8%, range [80.9, 97.1]). Sixth, we discarded pairs of epochs in which the Replay epoch contained missing samples, for instance, due to blinks or poor calibration that led to the temporary loss of the eye-tracking signal. This occurred in on average 5.4% (range [4.3, 7.4]) of epochs. Once data cleaning was completed, a median of 12423 pairs of epochs per observer (range [8311, 14026]) were analyzed.

#### Hypothesis tests

For hypothesis testing we used type-2 repeated-measures ANOVAs, as implemented in the ‘ez’ R package (Lawrence, 2016), with Greenhouse-Geisser p-value corrections for sphericity (*p*_*GG*_). Linear mixed-effects models were fitted with the ‘lme4’ R package (Bates, Mächler, Bolker, & Walker, 2015), using correlated random intercepts and slopes, to predict lambda-response latency and amplitude. Two-degree polynomial contrasts were used to capture the nonlinearity of the relationship between saccade size and response amplitude (Ries et al., 2018). Continuous fixed effects (e.g., saccade duration) were mean-centered and discrete fixed effects (e.g., viewing condition) were sum-coded (-0.5 vs 0.5). Confidence intervals (95% CIs) for estimated weights were approximated using parametric bootstrapping (2000 simulations). Model comparisons were performed based on the Bayes information criterion (BIC). Saccade duration, and not saccade amplitude or saccadic peak velocity, was used as the primary predictor because duration-based models fitted dependent variables decisively better than amplitude-based or velocity-based models. This was the case for response latency (offset-locked: BIC_amp_ = 1292.7, BIC_vel_ = 1186.9, BIC_dur_ = 1069.6; onset-locked: BIC_amp_ = 1153.2, BIC_vel_ = 1032.8, BIC_dur_ = 993.8) and, with one (explainable) exception, also response amplitude (offset-locked: BIC_amp_ = 795.7, BIC_vel_ = 677.1, BIC_dur_ = 630.4; onset-locked: BIC_amp_ = 717.0, BIC_vel_ = 696.7, BIC_dur_ = 704.1). To corroborate hypothesis tests based on CIs in cases in which evidence seemed weak, additional hierarchical model comparisons were performed based on the Bayes factor (BF) which was computed according to

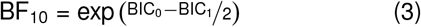

where BIC_0_ and BIC_1_ are the BIC of the null model (i.e., without the variable in question) and the alternative model (i.e., including the variable in question), respectively. In this formulation, BF_10_ encodes the likelihood by which the data would occur assuming the alternative model, as compared to assuming the null model. Nonlinear mixed-effects modeling was performed using the ‘nlme’ package (Pinheiro, Bates, DebRoy, Sarkar, & R Core Team, 2020). All error bars represent *±* one standard error of the mean, unless specified otherwise.

#### Computational modeling

Early visual responses to the consequences of real and simulated saccades were estimated using the model described in detail by Schweitzer, Seel, et al. (2025) with identical processing steps and parameter settings. We defined five spatial-frequency (SF) channels, that is, 0.25, 0.5, 1.0, 2.0, 4.0 cycles per degree of visual angle (cpd), and eight orientation channels separated by ^*π*^*/*_8_, from which a total of 40 log-Gabor filters and five SF-dependent temporal response functions (TRFs) were generated. TRFs were derived from predictions of Kelly’s spatiotemporal sensitivity function (Kelly, 1979, see also Figure 3B)

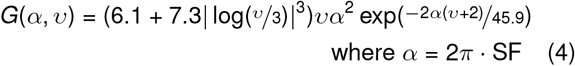

and approximated it accurately (*υ* represents velocity in dva/s; for details, see Schweitzer, Seel, et al., 2025). The TRF employed specifically in Figure 3E was adapted from previous work where it had been derived by fitting a gamma function to macaque V1 simple-cell spike-count data (Schweitzer, Doering, et al., 2025, their Eq. 10). The gamma distribution has been extensively used to simulate the visual system’s temporal-response characteristics (e.g., Groen et al., 2022; Teichert, Klingenhoefer, Wachtler, & Bremmer, 2010) and, as the derived TRF is monophasic, created a lowpass filter in the frequency domain.

Model simulations were run on 40 randomly selected scenes used in the experiment (with equal proportions of scenes with sharp and smooth edges, see Stimuli), converted to grayscale (RGB weights: [0.299, 0.587, 0.114]) applying the same gamma function as the presentation monitor (see Apparatus). Like during the experiment, scenes were placed within a simulated uniform screen with sharp borders and the same average luminance as the scene, whose simulated background was uniformly black (Figure 2A). The scene’s position relative to the simulated screen was constantly at center during simulated Saccade epochs and varied according the saccade trajectories (see below) during simulated Replay epochs. To reduce the computational complexity of the simulation, spatial resolution was downsampled to 11.2 pixels per degree and temporal resolution was set to 200 Hz.

Using the compressed exponential model (Han, Saunders, Woods, & Luo, 2013), that is,

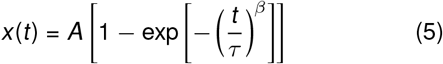

we created horizontal saccadic movement trajectories for six amplitudes *A* (1.5, 4.0, 6.5, 9.0, 11.5, and 14.0 dva; Figure 2B). Corresponding saccade durations (*D* = 2 · *τ*) were computed with Equation 2 using the median parameter estimates from individual fits of all observers’ main sequences (*k* = 20.1, range [14.8, 28.1]; *p* = 0.46, range [0.39, 0.53]), whereas the tailedness of saccades *β* was estimated from fitting Equation 5 to individual observers’ average saccade trajectories (*β* = 2.24, range [1.88, 2.53]). Saccadic peak-velocity estimates (Figure S4A), as well as when (relative to saccade onset) this peak velocity was reached (Figure S4B), were also extracted from these fits (Gibaldi & Sabatini, 2020). To simulate saccade curvature, we used the quadratic formula

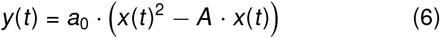

where *x*(*t*) is the horizontal saccade trajectory determined above, *A* its amplitude, and *a*_0_ is a scaling parameter set to ^0.1^*/*_*A*_. Saccade trajectories were padded with 350 ms pre-saccadic and 250 ms post-saccadic position samples. In the pre-saccadic interval the first 100 ms were used to linearly ramp-up the contrast of the simulated display, leading to a first pre-saccadic transient which was – due to the balanced alignment of saccade trajectories (see below) – identical across all tested amplitudes (Figure 2E). Note that we used a self-avoiding random-walk model (Engbert, Mergenthaler, Sinn, & Pikovsky, 2011) with additional smoothing of position estimates (Schweitzer & Rolfs, 2022) to simulate ocular drift movement (mean velocity: 1.3 dva/s; Figure 2B, right). This drift was added to the entire padded saccade trajectory.

To avoid any systematic contamination of model responses due to differences in start and landing positions of saccades with different amplitudes, simulated saccade trajectories were spatially aligned either to their start point (starting reference, constant pre-saccadic input) or to their landing point (landing reference, constant post-saccadic input), with either rightward or leftward direction (Figure 2A-B).

References were places at *±*8 dva horizontal eccentricity (and no vertical eccentricity) relative to the center of the simulated screen. Around each gaze-position sample – varying during Saccade trials and constant during Replay trials – an image with a size of 90 *×* 60 dva was bilinearly interpolated from the simulated screen (retinal view; Figure 2C), resulting in a x-y-t matrix with a size of 1011 *×* 674 *×* 134 that served as input to the model. In each simulation trial, model responses (Figure 2E-H) were summed across SF channels (except for analyses shown in Figure S5) and orientation channels, as well as across the spatial dimensions. Aggregated responses for each simulated condition (viewing condition *×* amplitude) were computed by collapsing across starting/landing references and rightward/leftward directions. Response amplitudes were denoted the maximum value of the waveform, whereas latencies were defined as the time point at which responses surpassed 1/3 of that amplitude (except for the profiles shown in Figure 3F where half-max latency was extracted).

## Acknowledgments

We thank Melkeberhan Degefa for her indispensable help with data collection, as well as Massimo Vescovi and Mauro Zago for their technical support.

## Author contributions

Conceptualization: RS, CHH

Methodology: RS

Funding acquisition: CHH

Project administration: CHH, RS

Investigation: RS

Data curation: RS

Formal analysis: RS

Visualization: RS

Software: RS

Supervision: CHH, OD

Writing – original draft: RS

Writing – review & editing: RS, CHH, OD

## Data and materials availability

Analysis and modeling scripts needed to evaluate the conclusions in the paper are made publicly available at https://github.com/richardschweitzer/LAM1_analysis_R. The study’s pre-registration, materials, and code are hosted on OSF and can be found at https://osf.io/crh8y. Raw data have been deposited in additional components of this repository: https://osf.io/rq2vc (Part 1), https://osf.io/7cdxj (Part 2), and https://osf.io/eygts (Part 3).

## Appendix A

### Supplementary Text

#### Evaluation of stimulus timing

To ensure that stimuli were presented correctly, we performed extensive timing tests with two photodiodes and DB-25 (parallel port) connector wired to an Arduino Due which sampled analog input at 1000 Hz. We devised a test in which a flickering grating, that is, with alternating low and high polarities, was presented with upon each screen refresh, based on which we were able to make four critical assertions. First, we ascertained the correct setup of the variable screen refresh rate set to 480 Hz. In a sample of 37,890 refreshes, the inter-flip interval (IFI) amounted to on average 2.12 ms (range [2.10, 2.34]). In other words, we achieved an effective frame rate of 472 Hz, notably without “dropping” frames – which would occur with classic fixed refresh rates and cause at least a doubling of the IFI. Second, to verify Gsync functionality, we introduced additional wait times (1.04–6.25 ms in steps of 1.04 ms) between stimulus rendering and flip. As wait times exceeded the nominal frame duration (i.e., 2.08 ms) IFIs increased linearly with wait times (*β* = 1.00, *t* = 1251.3, *p <* .001) while a constant delay of 0.43 ms (*β* = 0.43, *t* = 122.8, *p <* .001) incurred, likely due to stimulus preparation and drawing (Figure S1A). Clearly, neither slope (*β* = 0.001, *t* = 0.96, *p* = .338) nor intercept (*β* = −0.002, *t* = 0.59, *p* = .553) were affected by stimulus polarity. This behavior is expected from G-sync’s capability of flipping without having to wait for the monitor’s vertical retrace (Poth et al., 2018). Third, as shown in Figure S1B, photodiode responses clearly scaled with the administered wait times, showing that IFIs determined by the timestamps returned by the ‘flip’ command corresponded to physical display durations, thereby underlining the feasibility of OLED monitors for vision research (see also Dimigen, Badea, Simon, & Span, 2025). Fourth, we determined the video delay of the system, which should amount to approximately one refresh cycle at a frame rate of 480 Hz, that is, 2.08 ms. Indeed, a cross-correlation analysis of the trigger (‘off’ for low polarities and ‘on’ for high polarities) and photodiode signals revealed a maximal correlation at 2 ms (Figure S1C), providing good evidence that triggers and physical displays were closely aligned when the video delay was taken into account.

#### SF-specific model-response dynamics

Our modeling approach allowed looking at the saccade onset-locked latency dynamics of individual spatial frequency (SF) channels (Figure S5A). Model response latencies in channels selective to 0.25 cycles per degree of visual angle (cpd) hardly increased with increasing saccade duration (reference condition; 0.25 cpd: *β* = 0.10, *t* = 1.75, *p* = .096), but slopes significantly increased at higher SFs (treatment-coded difference to reference; 0.5 cpd: *β* = 0.20, *t* = 2.46, *p* = .023; 1.0 cpd: *β* = 0.71, *t* = 8.69, *p <* .001; 2.0 cpd: *β* = 0.82, *t* = 10.10, *p <* .001; 4.0 cpd: *β* = 0.76, *t* = 9.33, *p <* .001). As a comparison, measured lambda response latencies time-locked to saccade onset increased with saccade duration at a slope of 0.36 (cf. Figure 1G). Note that, for saccade-onset locked signals, slopes of 0 represent a locking of lambda responses to saccade onset, whereas slopes of 1 represent a locking to saccade offset. The results thus suggest that positive model responses of lower SF channels were driven by saccade-related visual events, whereas channels selective to higher SF were driven by fixation-related visual events. Normalized waveforms shown in Figure S5B confirm this notion: The low-SF (i.e., 0.25-cpd) channel showed saccade onset-locked increases in activity, while activity decreased in the high-SF (i.e., 2.0-cpd) channel. Moreover, post-saccadic activity increases were only observed in the high-SF channel where their onset latency clearly scaled with saccade duration. From these results it becomes evident that SF bands were variably affected by the same saccade-induced visual shift.

## Appendix B

### Supplementary Figures

**Figure S1.**
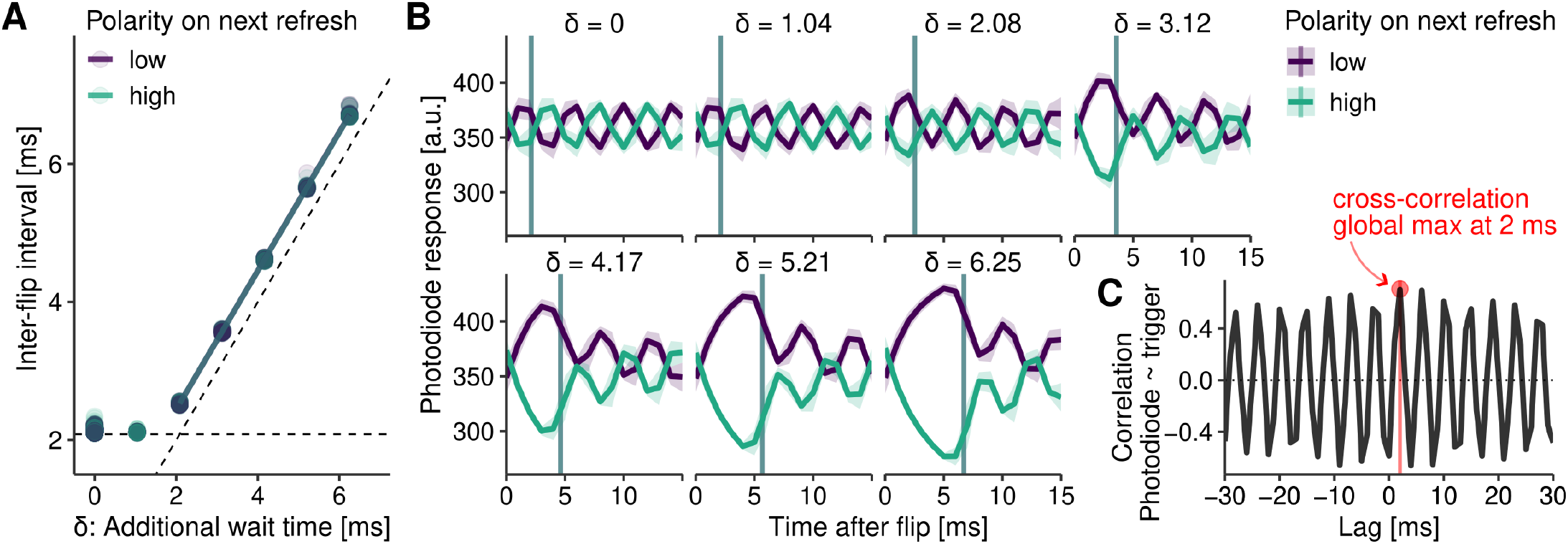
Evaluation of stimulus display timing. **A** Inter-flip intervals (IFI) during the presentation of binary stimulus flicker (low and high luminance) when additional wait times were inserted between stimulus rendering and flip. The horizontal dashed line indicates the minimal possible IFI at 480 fps. **B** Analog photodiode responses relative to the time of the flip. Individual panels correspond to instructed delay times (δ), whereas vertical lines represent the empirical IFI measured in this condition. **C** Cross-correlation of trigger and photodiode signals. The red dot indicates the maximum correlation (ρ = 0.70) found at a lag of 2 ms.

**Figure S2.**
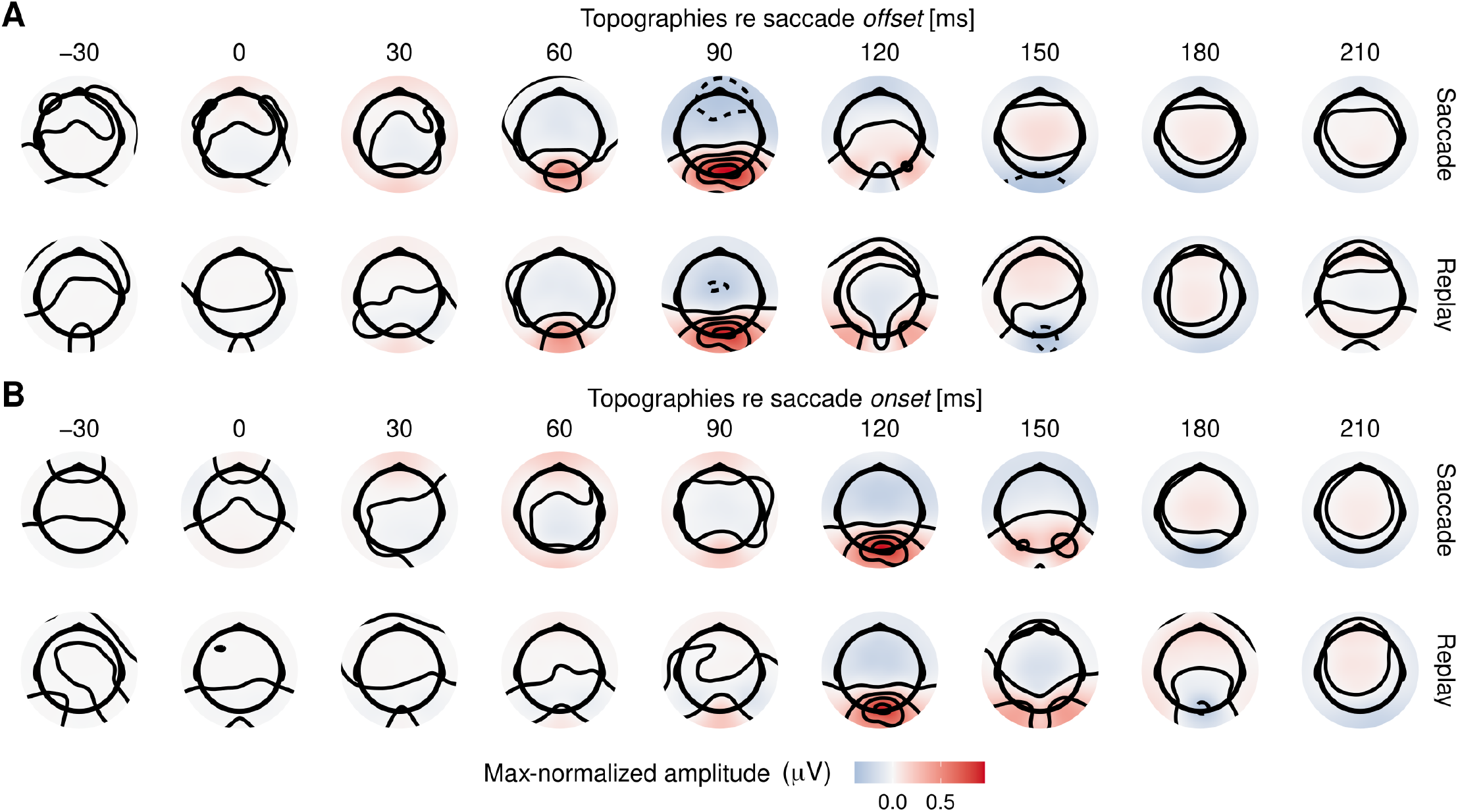
Topographies averaged across observers. For improved comparability topographies were normalized by dividing the maximum measured value across channels and time points relative to saccade offset (panel **A**) and onset (panel **B**) and Saccade and Replay condition (top and bottom rows, respectively).

**Figure S3.**
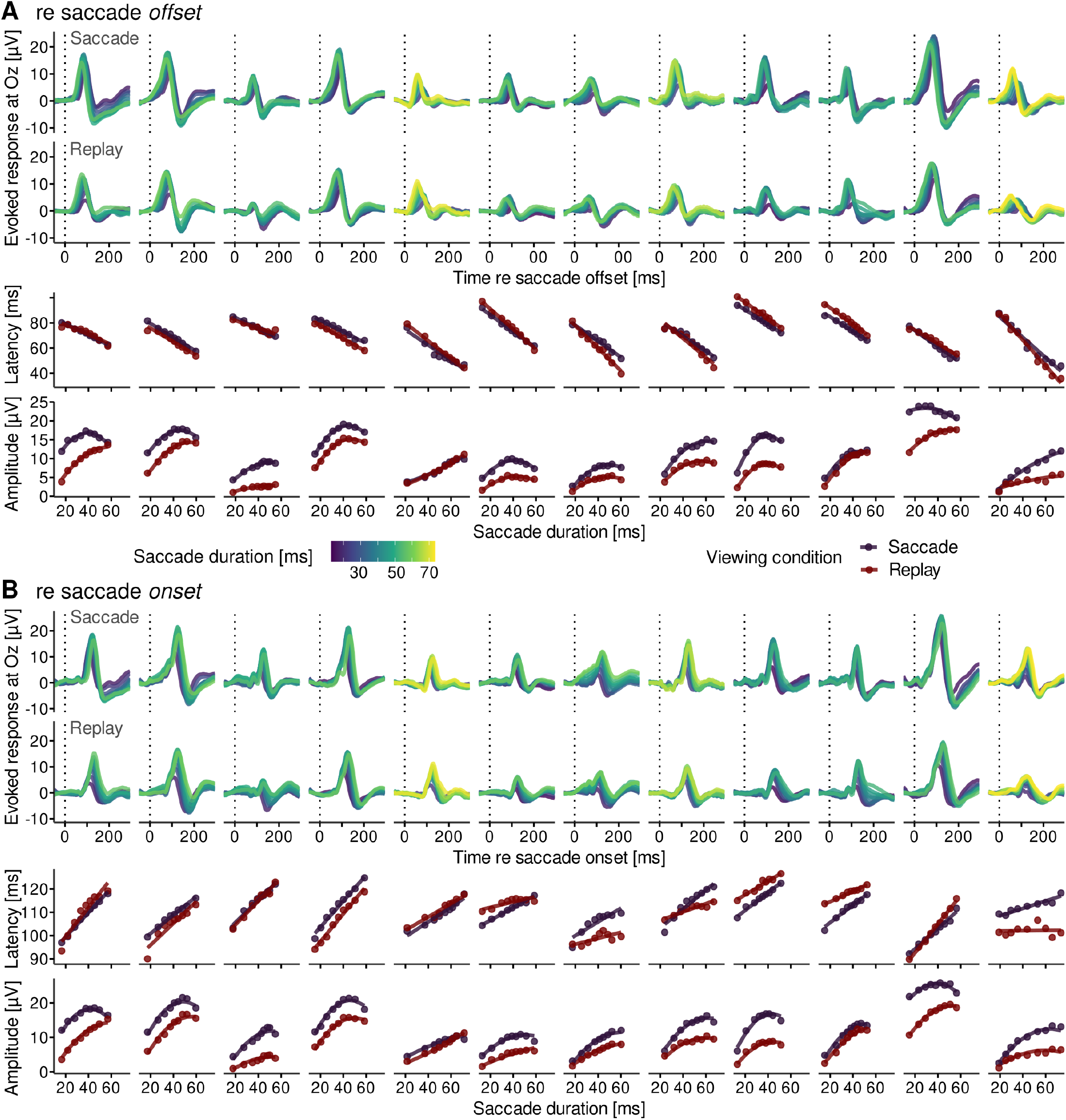
lambda responses in individual observers. **A** The first two rows show individual observers’ lambda waveforms time-locked to saccade offset in Saccade and Replay conditions with the same conventions as in Figure 1F. The third and fourth rows show extracted latency and amplitude data points for each viewing condition along with the predictions of mixed-effects modeling, again with the same conventions as in Figure 1G-H. Each column represents one observer. **B** Same as panel A, but locked to saccade onset.

**Figure S4.**
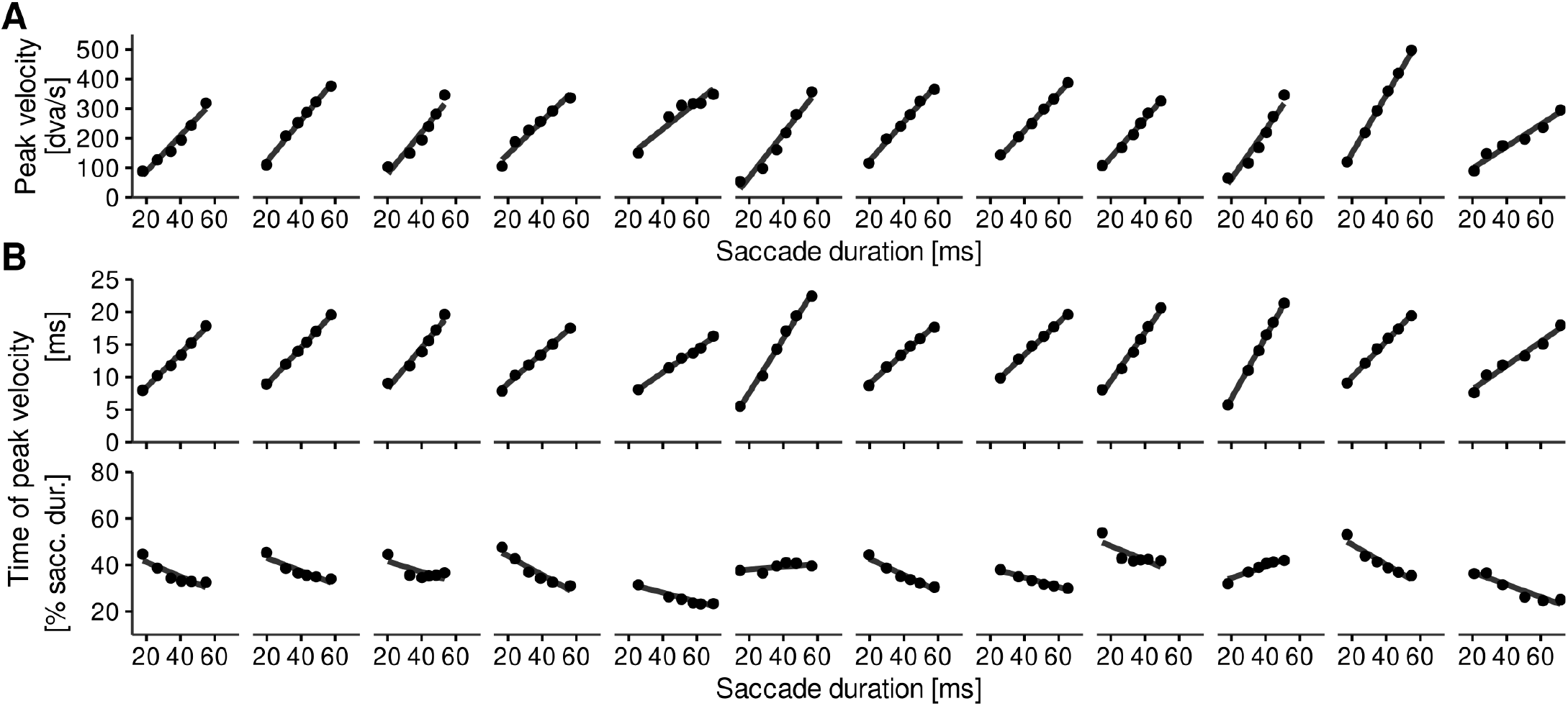
Individual observers’ saccadic peak-velocity dynamics. **A** Saccadic peak velocity, extracted from fitting the compressed exponential model (Equation 5) to averaged saccade trajectories (six bins per observer), as a linear function of the saccade’s average duration (β_0_ = 236.28, SE_0_ = 12.07, β_dur_ = 6.57, SE_dur_ = 0.51). Each column represents one observer. **B**) Same as above, but showing the time (as measured from saccade onset) when the peak velocity is reached, expressed as a linear function of saccade duration in absolute terms (top row; β_0_ = 13.94, SE_0_ = 0.28, β_dur_ = 0.29, SE_dur_ = 0.03) and as percentage of the saccade’s duration (bottom row; β_0_ = 36.34, SE_0_ = 1.43, β_dur_ = − 0.22, SE_dur_ = 0.06).

**Figure S5.**
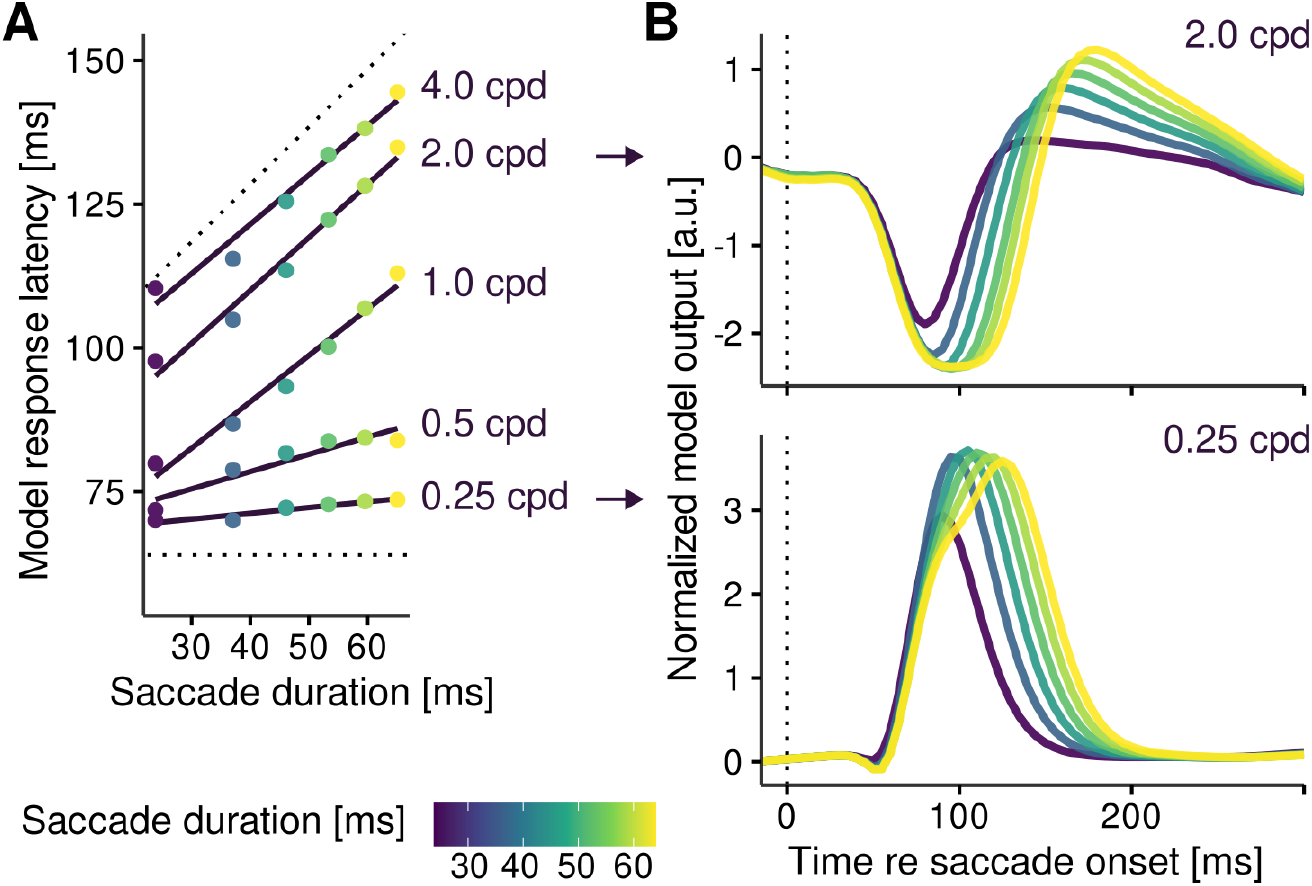
Model-response dynamics in individual spatial-frequency (SF) channels. **A** Model response latencies extracted from the responses of individual SF channels (averaged across orientation channels), as a function of saccade duration. Dotted lines represent hypothetical latency predictions that would occur if responses were locked to either saccade onset (β = 0) or saccade offset (β = 1). Annotations indicate the spatial frequency of the channel. **B** Average response waveforms of SF channels selective to 2.0 and 0.25 cycles per degree of visual angle (cpd).

